# How Fast Fast-Folding Proteins Fold in Silico

**DOI:** 10.1101/088849

**Authors:** Yuan-Ping Pang

**Affiliations:** Computer-Aided Molecular Design Laboratory, Mayo Clinic, Rochester, MN 55905, USA

**Keywords:** folding kinetics, folding rate, folding time, protein folding, protein simulation, molecular dynamics, survival analysis, FF12MC, CLN025, Trp-cage

## Abstract

In reported microcanonical molecular dynamics simulations, fast-folding proteins CLN025 and Trp-cage autonomously folded to experimentally determined native conformations. However, the folding times of these proteins derived from the simulations were more than 4–10 times longer than their experimental values. This article reports autonomous folding of CLN025 and Trp-cage in isobaric–isothermal molecular dynamics simulations with agreements within factors of 0.69–1.75 between simulated and experimental folding times at different temperatures. These results show that CLN025 and Trp-cage can now autonomously fold in silico as fast as in experiments, and suggest that the accuracy of folding simulations for fast-folding proteins begins to overlap with the accuracy of folding experiments. This opens new prospects of developing computer algorithms that can predict both ensembles of conformations and their interconversion rates for a protein from its sequence for artificial intelligence on how and when a protein acts as a receiver, switch, and relay to facilitate various subcellular-to-tissue communications. Then the genetic information that encodes proteins can be better read in the context of intricate biological functions.

## 1. Introduction

How fast can fast-folding proteins autonomously fold in silico? This question is important because experimental folding times (τs) [1-3] are rigorous benchmarks for evaluating the accuracy of protein folding simulations. If accurate, such simulations offer not only insight into protein folding pathways and mechanisms [4-7] but also a means to determine ensembles of conformations and their interconversion rates for a protein, which are responsible for “proteins to act as receivers, switches, and relays and facilitate communication from the subcellular level through to the cell and tissue levels” [8]. Due to approximations in the empirical potential energy functions for the folding simulations, most simulated τs reported to date have been much longer than the corresponding experimental τs. For example, early molecular dynamics (MD) simulations of fast-folding proteins using a distributed computing implementation with implicit solvation yielded τs that were consistent with the corresponding experimental values if Cα root mean square deviation (CαRMSD) cutoffs of 2.5–3.0 Å or 3.622 Å (in combination with a set of secondary structure criteria) were used to identify conformations that constitute the native structural ensembles [9,10]. However, according to the reported sensitivities of the simulated τs to CαRMSD cutoffs [9,10], the τs would be considerably longer than the experimental values, if typical CαRMSD cutoffs of <2.0 Å were used. For another example, advanced microcanonical MD simulations predicted τs of fast-folding proteins CLN025 [11] and Trp-cage [12] to be 600 ns at 343 K and 14 μs at 335 K, respectively [13]. These τs are of high quality as the τs were derived from the microcanonical MD simulations that resulted in the most populated conformations of CLN025 and Trp-cage with CαRMSDs of 1.0 and 1.4 Å from the experimental native conformations, respectively [13]. However, because the experimental τs of the two proteins reportedly increase as temperature decreases [1,2], the simulated τs at 300 K are conceivably more than 4–10 times longer than the experimental τs. Therefore, how fast fast-folding proteins fold in silico equates to how accurate protein folding simulations can be. Most reported τs to date suggest that fast-folding proteins cannot autonomously fold in silico as fast as in experiments. This implies an accuracy gap between simulation and experiment for protein folding rate (1/τ) that is determined by folding mechanism or pathways [14].

To narrow the accuracy gap, a new protein simulation method was developed. This method uses uniformly scaled atomic masses to compress or expand MD simulation time for improving configurational sampling efficiency or temporal resolution [15-17]. Uniformly reducing all atomic masses of a simulation system by tenfold can compress the simulation time by a factor of 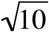 and hence improve the configurational sampling efficiency of the low-mass simulations at temperatures of ≤340 K [16]. As detailed in Refs. [15,16], this method facilitates protein folding simulations on personal computers (such as Apple Mac Pros) under isobaric–isothermal conditions at which most experimental folding studies are performed. As explained in Ref. [16], the kinetics of the low-mass simulation system can be converted to the kinetics of the standard-mass simulation system by simply scaling the low-mass time with a factor of 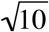. Subsequently, this low-mass simulation method led to the development of a revised AMBER forcefield that has shown improvements in (*i*) autonomously folding fast-folding proteins, (*ii*) simulating genuine localized disorders of folded globular proteins, and (*iii*) refining comparative models of monomeric globular proteins [18-20]. Hereafter the combination of the revised AMBER forcefield with the low-mass simulation method is termed FF12MC [18].

Further, in performing zebrafish toxicology experiments for a different project, this author observed that the times-to-death of the 20 toxin-treated fish varied widely in each experiment, although all 20 fish with nearly the same body weights received an intraperitoneal injection of the same dose of the same batch of botulinum neurotoxin serotype A. Yet, the mean time-to-death and its 95% confidence interval (95%CI) calculated using the open-source R survival package [21] varied slightly from one experiment to another. The resemblance of the live and dead states of the zebrafish to the unfolded and folded states of a protein inspired the use of the R survival package to predict τ of a fast-folding protein from its sequence as follows [16,18]: Perform (*i*) ≥20 distinct and independent MD simulations to autonomously fold a fast-folding protein sequence using FF12MC, which results in ≥20 sets of instantaneous protein conformations in time, (*ii*) a cluster analysis of all instantaneous conformations from the ≥20 sets to obtain the average conformation of the largest cluster and use the average conformation as the predicted native conformation of the protein, and (*iii*) a survival analysis using the ≥20 sets of the instantaneous conformations in time and the predicted native conformation to determine the mean τ and its 95%CI. As exemplified in Refs. [16,18], one advantage of this survival analysis method is that the τ prediction does assume that the fast-folding protein must follow a two-state folding mechanism; another advantage is rigorous estimation of mean τ and 95%CI from ≥20 simulations that are relatively short so that a few of these simulations may not capture a folding event.

As demonstrated below, use of the methods and forcefield outlined above resulted in accurate prediction of τs for CLN025 and Trp-cage (TC10b) and an answer to the important question of how fast fast-folding proteins fold in silico. A total of 160 distinct, independent, unrestricted, unbiased, isobaric–isothermal, microsecond MD simulations with a total aggregated simulation time of 1,011.2 μs were used for the prediction. All simulation times described hereafter have been converted to standard-mass simulation times.

## 2. Methods

### 2.1. Molecular dynamics simulations

A fast-folding protein in a fully extended backbone conformation was solvated with the TIP3P water [22] with surrounding counter ions and/or NaCls and then energy-minimized for 100 cycles of steepest-descent minimization followed by 900 cycles of conjugate-gradient minimization to remove close van der Waals contacts using SANDER of AMBER 11 (University of California, San Francisco). The resulting system was heated from 5 K to a temperature of 280–300 K at a rate of 10 K/ps under constant temperature and constant volume, then equilibrated for 10^6^ timesteps under constant temperature and constant pressure of 1 atm employing isotropic molecule-based scaling, and finally simulated in 40 distinct, independent, unrestricted, unbiased, and isobaric–isothermal MD simulations using PMEMD of AMBER 11 with a periodic boundary condition at 280–300 K and 1 atm. The fully extended backbone conformations (*viz.*, anti-parallel β-strand conformations) were generated by MacPyMOL Version 1.5.0 (Schrödinger LLC, Portland, OR). The numbers of TIP3P waters and surrounding ions, initial solvation box size, and ionizable residues are provided in Table S1. The 40 unique seed numbers for initial velocities of Simulations 1–40 are listed in Table S2. All simulations used (*i*) a dielectric constant of 1.0, (*ii*) the Berendsen coupling algorithm [23], (*iii*) the Particle Mesh Ewald method to calculate electrostatic interactions of two atoms at a separation of >8 Å [24], (*iv*) Δ*t* = 1.00 fs of the standard-mass time [18], (*v*) the SHAKE-bond-length constraints applied to all bonds involving hydrogen, (*vi*) a protocol to save the image closest to the middle of the “primary box” to the restart and trajectory files, (*vii*) a formatted restart file, (*viii*) the revised alkali and halide ions parameters [25], (*ix*) a cutoff of 8.0 Å for nonbonded interactions, (*x*) the atomic masses of the entire simulation system (both solute and solvent) were reduced uniformly by tenfold, and (*xi*) default values of all other inputs of the PMEMD module. The forcefield parameters of FF12MC are available in the Supporting Information of Ref. [16]. All simulations were performed on an in-house cluster of 100 12-core Apple Mac Pros with Intel Westmere (2.40/2.93 GHz).

### 2.2. Folding time estimation

The τ of a fast-folding protein was estimated from the mean time-to-folding in 40 distinct, independent, unrestricted, unbiased, and isobaric–isothermal MD simulations using survival analysis methods [21] implemented in the R survival package Version 2.38-3 (http://cran.r-project.org/package=survival). A Cα and Cβ root mean square deviation (CαβRMSD) cutoff of 0.98 Å was used to identify conformations that constitute the native structural ensemble. For each simulation with conformations saved at every 10^5^ timesteps, the first time-instant at which CαβRMSD reached ≤0.98 Å was recorded as an individual folding time (Table S3). Using the Kaplan-Meier estimator [26,27] [the Surv() function in the R survival package], the mean time-to-folding was first calculated from 40 simulations each of which captured a folding event at a low temperature of 280 K or 293 K. If a parametric survival function mostly fell within the 95%CI of the Kaplan-Meier estimation for these low-temperature simulations, the parametric survival function [the Surreg() function in the R survival package] was then used to calculate (*i*) the mean time-to-folding of the 40 low-temperature simulations and (*ii*) the mean time-to-folding of 40 new simulations, which were identical to the low-temperature simulations except that the temperature was increased to 300 K.

### 2.3. Cluster analysis and data processing

The conformational cluster analyses of CLN025 and TC10b were performed using CPPTRAJ of AmberTools 16 (University of California, San Francisco) with the average-linkage algorithm [28], epsilon of 2.0 Å, and root mean square coordinate deviation on all Cα and Cβ atoms (see Table S4). No energy minimization was performed on the average conformation of any cluster. The linear regression analysis was performed using the PRISM 5 program for Mac OS X, Version 5.0d (GraphPad Software, La Jolla, California).

## 3. Results and discussion

### 3.1. Simulated folding times of β-protein CLN025 at different temperatures

To determine how fast β-protein CLN025 autonomously folds in silico, 40 distinct, independent, unrestricted, unbiased, and isobaric–isothermal, and 3.16-μs MD simulations of CLN025 were performed at 300 K. A cluster analysis of these simulations revealed that the average conformation of the largest cluster adopted a β-hairpin conformation [Fig. 1(A)]. This average conformation had a CαRMSD of 0.87 Å and a CαβRMSD of 0.94 Å relative to the average conformation of 20 NMR-determined conformations of CLN025 [1] [Fig. 1(B)]. Using (*i*) the average conformation of the largest cluster as the predicted native conformation of CLN025 and (*ii*) CαβRMSD of ≤0.98 Å from the predicted native conformation to define the native structural ensemble of CLN025, the first time-instant at which CαβRMSD of the full-length CLN025 sequence reached ≤0.98 Å was recorded as an individual folding time for each of the 40 simulations (Table S3A). CαβRMSD was used here to determine the individual folding time because it has both main-chain and side-chain structural information and is more stringent to measure structural similarity than CαRMSD. Plotting the natural logarithm of the nonnative state population of CLN025 versus time-to-folding revealed a linear relationship with r^2^ of 0.97 (Fig. 2), which indicates that CLN025 follows a two-state folding mechanism. Using the 40 individual folding times as times-to-folding, a survival analysis predicted the τ of CLN025 to be 198 ns (95%CI = 146–270 ns; n = 40) at 300 K (Table 1). These results agree with the experimental finding that the folding of CLN025 followed a two-state folding mechanism with a τ of 137 ns at 300 K, where the τ was obtained from Fig. 6 of Ref. [1]. To substantiate the agreement between the experimental and simulated τs at 300 K, the 40 CLN025 simulations were repeated at 293 K. Using the same CαβRMSD cutoff and the same predicted native conformation, a survival analysis showed that CLN025 followed a two-state folding mechanism (r^2^ = 0.94; Fig. 2) with a τ of 279 ns (95%CI = 204–380 ns; n = 40) at 293 K (Table 1). This outcome again agrees with the experimental τ of 261 ns at 293 K that was also obtained from Fig. 6 of Ref. [1].

**Fig. 1.**
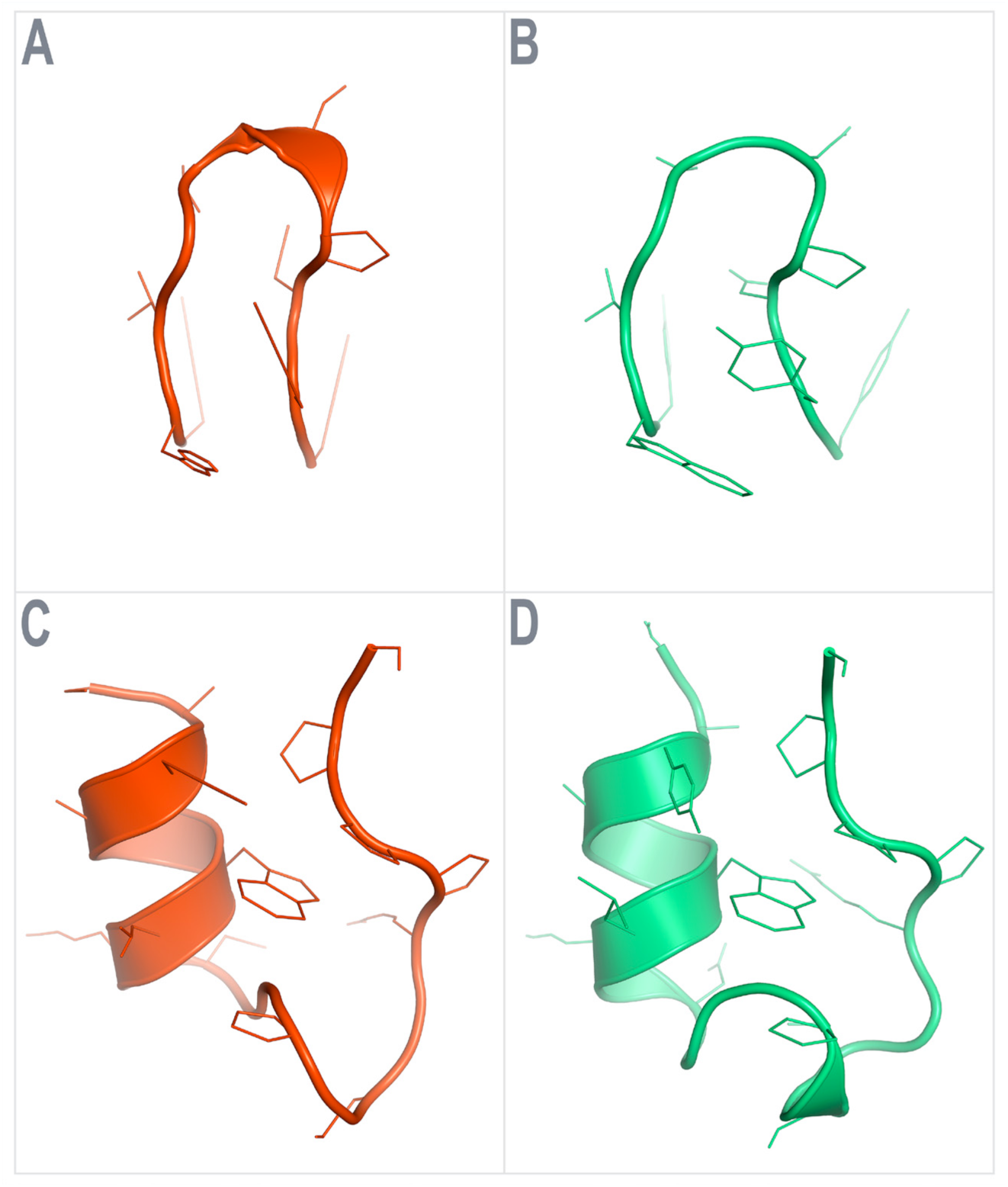
Native conformations of CLN025 and Trp-cage (TC10b) derived from experiments and simulations. (A) The average CLN025 conformation of the largest cluster of the 40 distinct, independent, unrestricted, unbiased, and isobaric–isothermal, and 3.16-μs MD simulations using FF12MC. (B) The average of 20 CLN025 NMR structures (PDB ID: 2RVD). (C) The average Trp-cage conformation of the largest cluster of the 40 distinct, independent, unrestricted, unbiased, and isobaric–isothermal, and 9.48-μs MD simulations using FF12MC. (D) The average of 28 Trp-cage NMR structures (PDB ID: 2JOF).

**Fig. 2.**
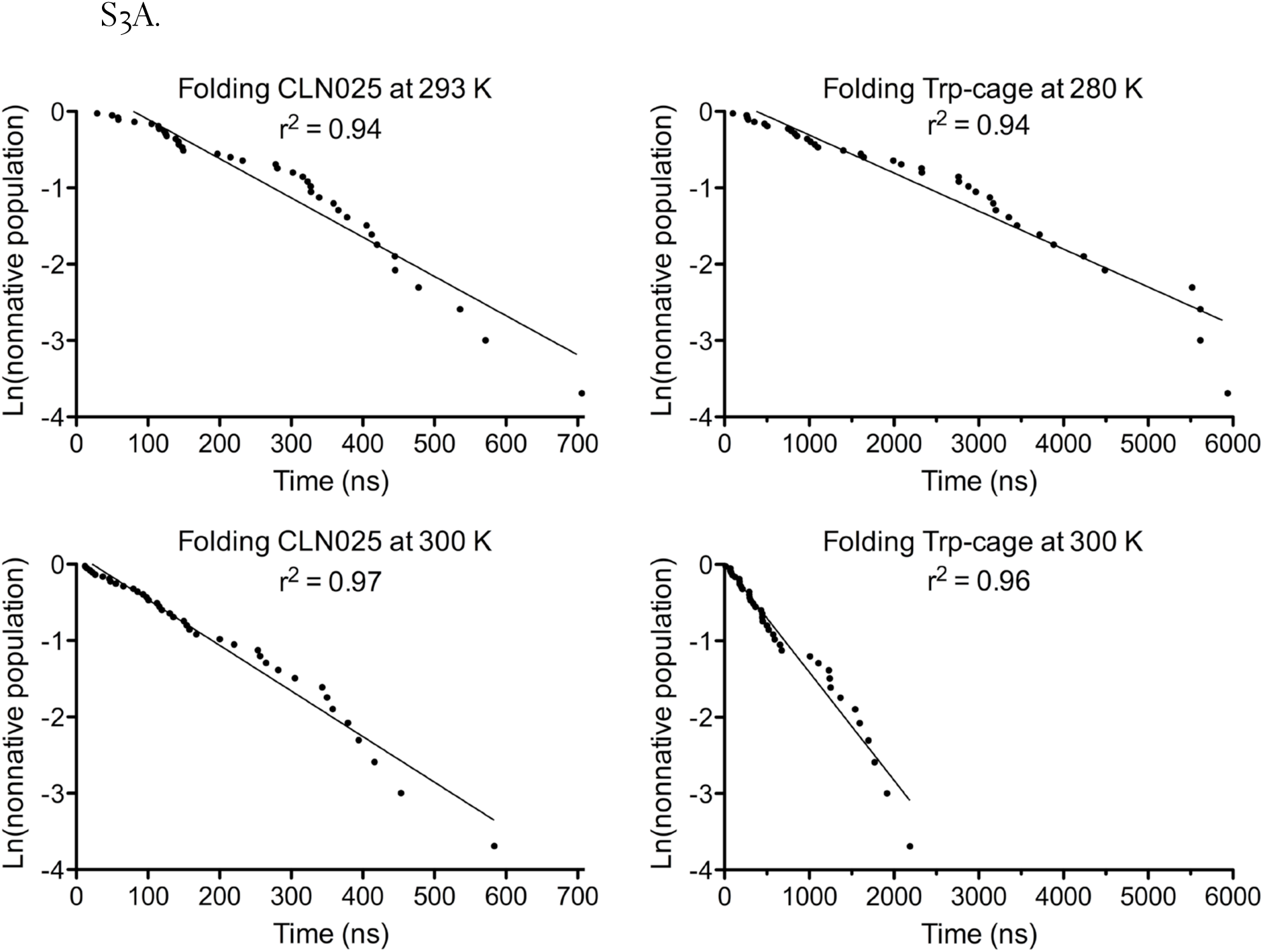
Plots of the natural logarithm of the nonnative state population of CLN025 and Trp-cage (TC10b) over time-to-folding. The individual folding times were taken from Table S3A.

**Table 1.**
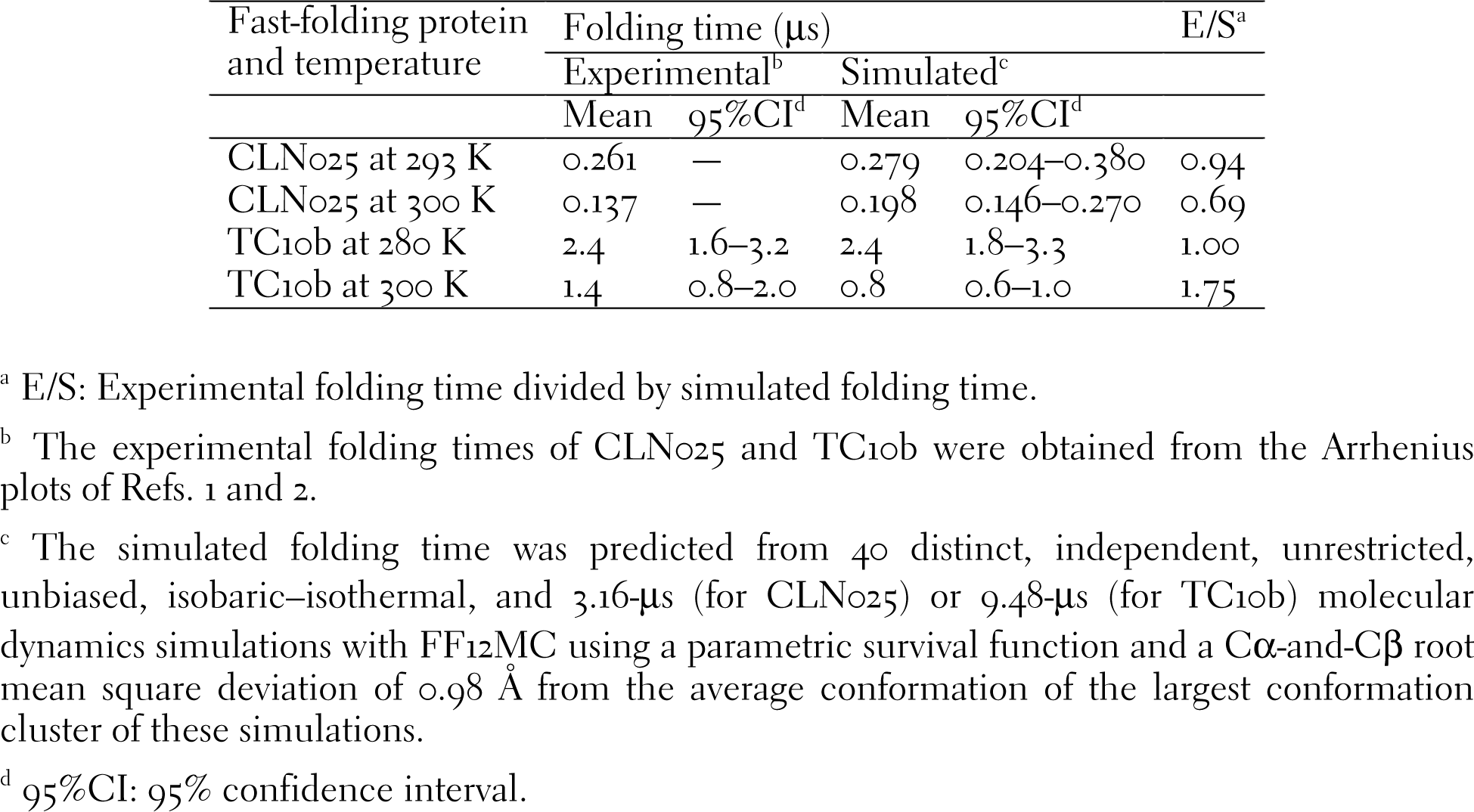
Experimental and simulated folding times of CLN025 and Trp-cage (TC10b)

### 3.2. Simulated folding times of α-protein Trp-cage at different temperatures

To determine how fast α-protein Trp-cage autonomously folds in silico, 40 distinct, independent, unrestricted, unbiased, and isobaric–isothermal, and 9.48-μs MD simulations of the Trp-cage (TC10b) sequence were performed at 280 K. The average conformation of the largest cluster of these simulations [Fig. 1(C)] had a CαRMSD of 1.69 Å and a CαβRMSD of 1.86 Å from the average conformation of 28 NMR-determined conformations of TC10b [12] [Fig. 1(D)]. Using the CαβRMSD cutoff of 0.98 Å and the average conformation of the largest cluster as the predicted native conformation, a survival analysis showed that TC10b followed a two-state folding mechanism (r^2^ = 0.94; Fig. 2) in the simulations with a τ of 2.4 μs (95%CI = 1.8–3.3 μs; n = 40) at 280 K (Table 1). These results are consistent with the experimentally determined two-state folding mechanism and the experimental τ of 2.4 μs (95%CI = 1.6–3.2 μs) at 280 K for Trp-cage. The experimental τ was extrapolated from Fig. 4 of NMR ln*k*_F_ in Ref. [2]. The experimental 95%CI was calculated from the reported errors of ±0.18 for ln*k*_F_ in the 12–14 range [2] according to the standard method for propagation of errors of precision [29]. Repeating the 40 simulations of TC10b at 300 K and the survival analysis using the same predicted native conformation and the CαβRMSD cutoff of 0.98 Å revealed a two-state folding mechanism (r^2^ =0.96; Fig. 2) and a simulated τ of 0.8 μs (95%CI = 0.6–1.0 μs; n = 40; Table 1), which is also consistent with the experimental τ of 1.4 μs (95%CI = 0.8–2.0 μs) at 300 K [2].

### 3.3. Convergence of the simulated folding times

The simulated τ may not be converged and useful if the number of simulations is insufficient or if a lax CαRMSD or CαβRMSD is used to define the native structural ensemble. To assess the convergence of the simulated τs of CLN025 and Trp-cage described above, the average conformations of the largest cluster (*viz.,* the predicted native conformations) for CLN025 and Trp-cage shown in Fig. 1 and the corresponding simulated τs listed in Table 1 were re-generated using Simulations 1–20 or 1–30, respectively. According to the CαβRMSDs in Table S4, the average conformation of the largest cluster of Simulations 1–40 for CLN025 is identical to the corresponding ones of Simulations 1–20 and Simulations 1–30, and the same is true for Trp-cage. Further, the resulting changes in the mean and 95%CI of the simulated τ are negligible when the number of simulations increased from 20 to 30 or 40 (Table S5). These results indicate that 40 simulations are sufficient.

In the present study, a stringent CαβRMSD of ≤0.98 Å for the full sequence of CLN025 or Trp-cage was used to define the native structural ensemble, in contrast to the use of CαRMSD for a fast-folding protein with truncations on terminal residues. However, using an “overly” stringent CαβRMSD cutoff of 0.98 Å may lengthen the simulated τ, whereas using the average rather than the representative instantaneous conformation of the largest cluster as the predicted native conformation may shorten the simulated τ. To address these concerns, all τs in Table 1 were re-estimated from the same simulations using both the average and representative conformations with CαβRMSD cutoffs that varied from 0.98 Å to 1.40 Å. As apparent from Table S6, the simulated τs of CLN025 and Trp-cage are insensitive to the change from the average to the representative conformation, and these τs are also insensitive to the variation of the CαβRMSD cutoff within 0.98–1.40 Å. In addition, all τs in Table 1 were determined from trajectories that revealed not only hazard functions of a two-state folding mechanism for both CLN025 and Trp-cage (Fig. 2) but also their consistent folding events, which are traceable in Videos S1 and S2 (videos of long folding events are not shown due to file size limit). The theoretical mechanism and folding events are consistent with their experimentally determined folding mechanism and native conformations. It is therefore conceivable that the simulated τs in Table 1 are converged and may be used to explain, confirm, or predict folding rates of CLN025 and Trp-cage.

### 3.4. Significance of the simulated folding times

The present study shows that agreements within factors of 0.69–1.75 between the experimental and simulated τs have been achieved for CLN025 and Trp-cage (Table 1). These agreements indicate that fast-folding proteins CLN025 and Trp-cage can now autonomously fold in simulations as fast as in experiments, and provide an answer to the important question of how fast fast-folding proteins fold in silico. These agreements also suggest that the accuracy of folding simulations for fast-folding proteins is beginning to overlap with the accuracy of folding experiments. This opens new prospects of combining simulation with experiment to develop computer algorithms that can predict ensembles of conformations and their interconversion rates for a protein from its sequence. Such algorithms can improve the artificial intelligence on how and when “proteins act as receivers, switches, and relays and facilitate communication from the subcellular level through to the cell and tissue levels” [8]. Then the genetic information that encodes proteins can be better read in the context of intricate biological functions.

## Acknowledgments

The author acknowledges the support of this work from the US Defense Advanced Research Projects Agency (DAAD19-01-1-0322), the US Army Medical Research and Material Command (W81XWH-04-2-0001), the US Army Research Office (DAAD19-03-1-0318, W911NF-09-1-0095, and W911NF-16-1-0264), the US Department of Defense High Performance Computing Modernization Office, and the Mayo Foundation for Medical Education and Research. The contents of this article are the sole responsibility of the author and do not necessarily represent the official views of the funders.

## Supplementary Information

Tables S1–S6 and Videos S1 and S2

## References

[1] C.M. Davis, S.F. Xiao, D.P. Raeigh, R.B. Dyer, Raising the speed limit for β-hairpin formation, J. Am. Chem. Soc. 134 (2012) 14476–14482.

[2] A. Byrne, D.V. Williams, B. Barua, S.J. Hagen, B.L. Kier, N.H. Andersen, Folding dynamics and pathways of the trp-cage miniproteins, Biochemistry 53 (2014) 6011–6021.

[3] J. Kubelka, J. Hofrichter, W.A. Eaton, The protein folding 'speed limit', Curr Opin Struct Biol 14 (2004) 76–88.

[4] B. Gillespie, K.W. Plaxco, Using protein folding rates to test protein folding theories, Annu. Rev. Biochem. 73 (2004) 837–859.

[5] C.D. Snow, E.J. Sorin, Y.M. Rhee, V.S. Pande, How well can simulation predict protein folding kinetics and thermodynamics?, Annu. Rev. Biophys. Biomol. Struct. 34 (2005) 43–69.

[6] H. Gelman, M. Gruebele, Fast protein folding kinetics, Q Rev Biophys 47 (2014) 95–142.

[7] V. Munoz, M. Cerminara, When fast is better: protein folding fundamentals and mechanisms from ultrafast approaches, Biochem. J. 473 (2016) 2545–2559.

[8] V.J. Vinson, Proteins in motion. Introduction, Science 324 (2009) 197.

[9] C.D. Snow, N. Nguyen, V.S. Pande, M. Gruebele, Absolute comparison of simulated and experimental protein-folding dynamics, Nature 420 (2002) 102–106.

[10] C.D. Snow, B. Zagrovic, V.S. Pande, The Trp cage: folding kinetics and unfolded state topology via molecular dynamics simulations, J. Am. Chem. Soc. 124 (2002) 14548–14549.

[11] S. Honda, T. Akiba, Y.S. Kato, Y. Sawada, M. Sekijima, M. Ishimura, A. Ooishi, H. Watanabe, T. Odahara, K. Harata, Crystal structure of a ten-amino acid protein, J. Am. Chem. Soc. 130 (2008) 15327–15331.

[12] B. Barua, J.C. Lin, V.D. Williams, P. Kummler, J.W. Neidigh, N.H. Andersen, The Trp-cage: Optimizing the stability of a globular miniprotein, Protein Eng. Des. Sel. 21 (2008) 171–185.

[13] K. Lindorff-Larsen, S. Piana, R.O. Dror, D.E. Shaw, How fast-folding proteins fold, Science 334 (2011) 517–520.

[14] A.R. Fersht, On the simulation of protein folding by short time scale molecular dynamics and distributed computing, Proc. Natl. Acad. Sci. U.S.A. 99 (2002) 14122–14125.

[15] Y.-P. Pang, Low-mass molecular dynamics simulation: A simple and generic technique to enhance configurational sampling, Biochem. Biophys. Res. Commun. 452 (2014) 588–592.

[16] Y.-P. Pang, Low-mass molecular dynamics simulation for configurational sampling enhancement: More evidence and theoretical explanation, Biochem. Biophys. Rep. 4 (2015) 126–133.

[17] Y.-P. Pang, Use of multiple picosecond high-mass molecular dynamics simulations to predict crystallographic B-factors of folded globular proteins, Heliyon 2 (2016) e00161.

[18] Y.-P. Pang, FF12MC: A revised AMBER forcefield and new protein simulation protocol, Proteins 84 (2016) 1490–1516.

[19] Y.-P. Pang, At least 10% shorter C–H bonds in cryogenic protein crystal structures than in current AMBER forcefields, Biochem. Biophys. Res. Commun. 458 (2015) 352–355.

[20] Y.-P. Pang, Use of 1–4 interaction scaling factors to control the conformational equilibrium between α-helix and β-strand, Biochem. Biophys. Res. Commun. 457 (2015) 183–186.

[21] T.M. Therneau, P.M. Grambsch, Modeling Survival Data: Extending the Cox Model, Springer-Verlag, New York, 2000.

[22] W.L. Jorgensen, J. Chandreskhar, J.D. Madura, R.W. Impey, M.L. Klein, Comparison of simple potential functions for simulating liquid water, J. Chem. Phys. 79 (1983) 926–935.

[23] H.J.C. Berendsen, J.P.M. Postma, W.F. van Gunsteren, A. Di Nola, J.R. Haak, Molecular dynamics with coupling to an external bath, J. Chem. Phys. 81 (1984) 3684–3690.

[24] T.A. Darden, D.M. York, L.G. Pedersen, Particle mesh Ewald: An N log(N) method for Ewald sums in large systems, J. Chem. Phys. 98 (1993) 10089–10092.

[25] I.S. Joung, T.E. Cheatham, Determination of alkali and halide monovalent ion parameters for use in explicitly solvated biomolecular simulations, J. Phys. Chem. B 112 (2008) 9020–9041.

[26] E.L. Kaplan, P. Meier, Nonparametric estimation from incomplete observations, J. Am. Stat. Assoc. 53 (1958) 457–481.

[27] J.T. Rich, J.G. Neely, R.C. Paniello, C.C. Voelker, B. Nussenbaum, E.W. Wang, A practical guide to understanding Kaplan-Meier curves, Otolaryngol. Head Neck Surg. 143 (2010) 331–336.

[28] J. Shao, S.W. Tanner, N. Thompson, T.E. Cheatham III, Clustering molecular dynamics trajectories: 1. Characterizing the performance of different clustering algorithms, J. Chem. Theory Comput. 3 (2007) 2312–2334.

[29] J. Andraos, On the propagation of statistical errors for a function of several variables, J. Chem. Educ. 73 (1996) 150–154.

